# Deep Proteome Profiling of Metabolic Dysfunction-Associated Steatotic Liver Disease

**DOI:** 10.1101/2024.05.24.595658

**Authors:** Felix Boel, Vyacheslav Akimov, Mathias Teuchler, Mike Krogh Terkelsen, Charlotte Wilhelmina Wernberg, Frederik Tibert Larsen, Philip Hallenborg, Mette Munk Lauridsen, Aleksander Krag, Susanne Mandrup, Kim Ravnskjær, Blagoy Blagoev

## Abstract

Metabolic dysfunction-associated steatotic liver disease (MASLD) affects roughly 1 in 3 adults and is a leading cause of liver transplants and liver related mortality. A deeper understanding of disease pathogenesis is essential to assist in developing blood-based biomarkers. Here, we use data-independent acquisition mass spectrometry to assess disease-state associated protein profiles in human liver, blood plasma, and white adipose tissue (WAT). In liver, we find that MASLD is associated with an increased abundance of proteins involved in immune response and extracellular matrix (ECM) and a decrease in proteins involved in metabolism. Cell type deconvolution of the proteome indicate liver endothelial and hepatic stellate cells as main source of the ECM rearrangements, and hepatocytes as the major contributor to the changes in liver metabolism. In the blood, profiles of several MASLD-associated proteins that correlate with their expression in WAT rather than liver yet could serve as suitable liver disease predictors in a multi-protein panel marker. Moreover, our proteomics-based logistic regression models consistently outperform existing methods for predicting MASLD and liver fibrosis from human blood samples.

## Introduction

Advanced stages of steatotic liver disease (SLD) gradually impair liver function, resulting in end-stage non-reversible chronic liver disease (CLD)^1^. CLD causes more than 2 million deaths annually, and treatment requires liver transplantation. The most prevalent subtype of SLD, MASLD – formerly known as non-alcoholic fatty liver disease (NAFLD)^2^ – affects more than 30 % of all adults^3^. Early stages of MASLD are asymptomatic, making preventive screening and disease monitoring a difficult task. Currently, accurate diagnosis relies on a liver biopsy to histologically assess steatosis, lobular inflammation, hepatocellular ballooning. Based on the scoring, MASLD become categorized as metabolic dysfunction-associated steatotic liver (MASL) or metabolic dysfunction-associated steatohepatitis (MASH)^4^. In MASH, tissue repair processes lead to an accumulation of tissue scarring (fibrosis), giving rise to cirrhosis, CLD, and hepatocellular carcinoma (HCC)^5^.

Liver biopsies are invasive, non-standard, prone to sampling error, and at risk of intraobserver variation^6^. Consequently, many efforts have been made to utilize readily available biochemistry and clinical information to help grade disease progression and thereby reduce unnecessary biopsies. However, screenings are not standardized and vary depending on the country, research laboratory, and individual health professionals. Moreover, these methods, deriving from correlation-based studies, does not consider interplay with other metabolic disorders and factors of human heterogeneity such as age, gender, ethnicity, genetics, epigenetics, smoking, alcohol consumption, obesity, gut microbiota, and hormonal imbalances^7^. As a result, clinicians face a shortage of tests for precise diagnosis, prognosis, disease monitoring, prediction, and assessing the effectiveness of interventions.

Blood sampling is the preferred method in non-invasive diagnostics^8^, aiming to measure proteins that end up in circulation either by cell secretion or cell leakage. This makes proteomics an excellent tool in biomarker discovery^9^. To ensure proteins found in the blood are indeed disease-specific, consideration must be given to the biological changes in a tissue-specific context of the proteome^10^. However, no studies have taken a purely proteomics-based approach to explore MASLD pathogenesis collectively in human liver and blood samples.

Recent advances in speed and sensitivity of mass spectrometers have unlocked methods for data-independent acquisition (DIA) proteomics^11^. Compared to conventional data-dependent acquisition (DDA), DIA eliminates bias of high abundance peptides, decreases run-to-run variation, improves sample reproducibility, and decreases required input material, all of which are vital attributes when analyzing highly variable human samples from low starting material^12^. Furthermore, DIA avoids the bias towards high abundant peptides and can consistently quantify hundreds even in blood^13^. This makes it an exciting platform for discovery of novel blood biomarkers, and, with increasing instrument speed and automation efforts, this approach holds a potential for high-throughput screening of blood samples in a clinical setting.

In this study, we perform DIA-based deep proteomic profiling of samples acquired from patients at risk of MASLD with ≥class 2 obesity (BMI > 35). From liver biopsies, we do an explorative characterization of MASLD pathogenesis, including analysis of signaling pathways, cell type-specific responses, and transcriptional regulation. Additionally, we screen blood plasma for biomarkers of MASLD and highlight the importance of considering the effects of other tissues when investigating multifaceted diseases.

## Results

### Study design and data collection

Our cohort consisted of patients with a BMI > 35 (kg m^-2^) at risk of metabolic disorder. Most study participants were female, with a median age in the late 40s and a nearly 50% estimated body fat (Table 1). As part of the study protocol, all patients had a percutaneous suction needle liver biopsy, which was assessed by an experienced liver pathologist, using NAFLD activity score (NAS) and Kleiner fibrosis grade. To determine their MASLD status, we used the SAF algorithm, which divides patients into no-MASLD, MASL and MASH^14^. For proteomics, we utilized DIA-MS workflow^11^ (Fig. 1a) to analyze liver biopsies (n = 58), blood plasma (n = 143), scWAT biopsies (n = 58), and oWAT biopsies (n = 27). Additionally, as a validation cohort, we had 2-year follow-up blood plasma (n = 41) from a subset of the patients, some of whom had undergone bariatric surgery during follow-up. In liver, we quantified 7096 unique proteins with similar proteome coverage in each sample (Supplementary Fig. S2a,b). In scWAT and oWAT, we quantified 6945 and 6979 unique proteins respectively (Supplementary Fig. S2c,d). While in plasma, using short gradient DIA, we quantified 578 unique proteins (Supplementary Fig. S2e).

**Fig. 1:**
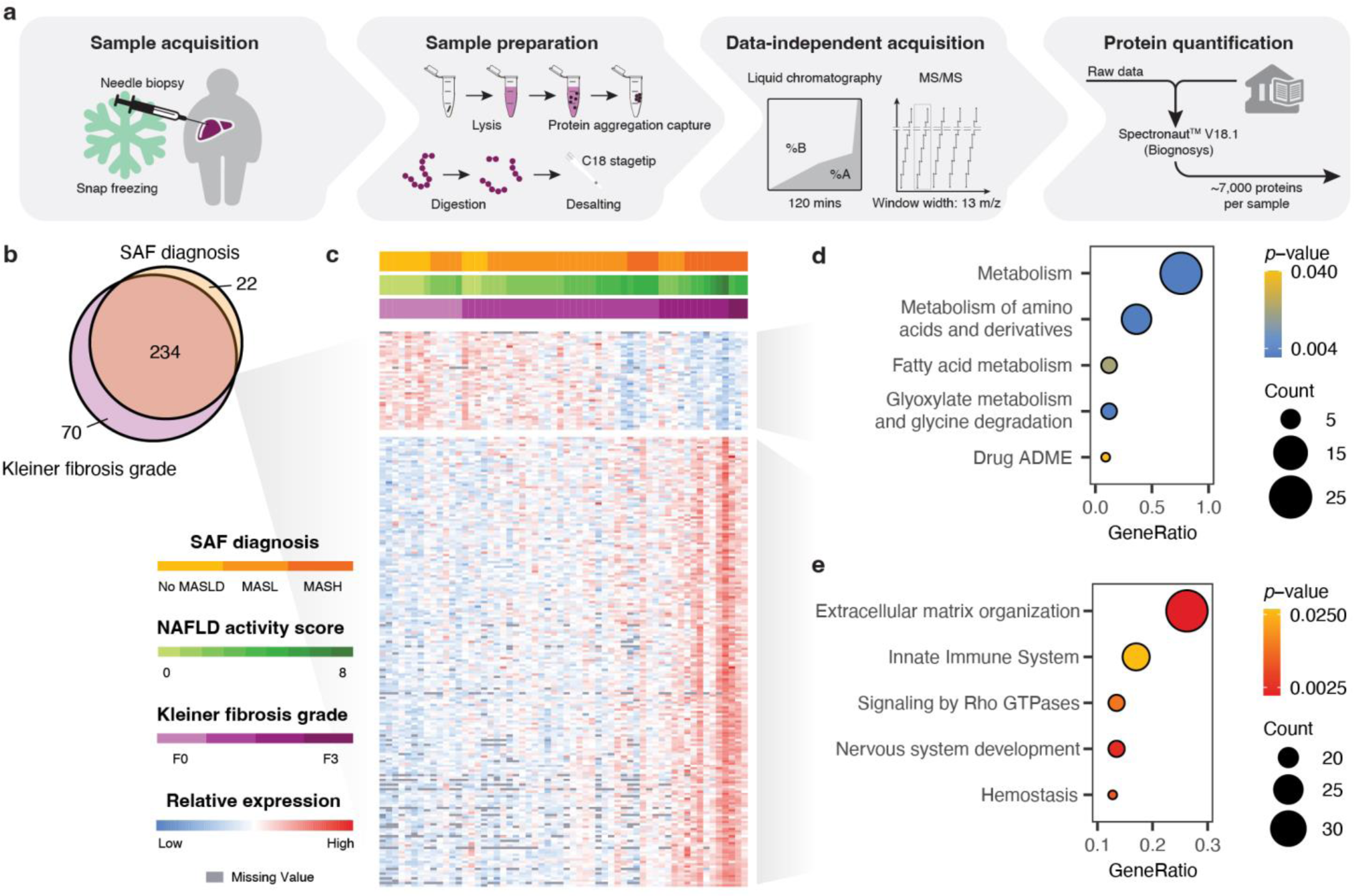
Identification of proteins related to MASLD pathogenesis. **a,** Workflow of our data acquisition pipeline for liver samples. **b,** Venn diagram of differentially expressed proteins for Kleiner fibrosis grade and SAF diagnosis. **c,** Heatmap for relative expression of the 234 differentially expressed proteins shared between Kleiner fibrosis grade and SAF diagnosis, shown as the relative expression across patients (columns). Rows (proteins) were sorted based on hierarchical clustering, while columns (patients) were sorted according to increasing (i) Kleiner fibrosis grade, (ii) SAF diagnose, and (iii) NAFLD activity score. **d,e,** Pathway enrichment analysis based on the Reactome pathway database for proteins upregulated (**d**) and downregulated (**e**) with increasing MASLD histopathology.

**Table 1:** Patient characteristics. Patient data for samples analyzed by proteomics shown as medians ± standard deviation (s.d.) or summarized counts.

| Clinical feature | Liver<br>(n = 58) | scWAT<br>(n = 58) | oWAT<br>(n = 27) | Plasma<br>(n = 143) | Plasma (follow-up)<br>(n = 41) |
| --- | --- | --- | --- | --- | --- |
| Gender (% female) | 65.5 | 65.5 | 74.1 | 78.3 | 75.6 |
| Age (years) | 47.5 ± (13.6) | 46.0 ± (13.5) | 45.0 ± (11.9) | 46.0 ± (12.3) | 53.0 ± (13.0) |
| BMI (kg m<sup>-2</sup>) | 42.6 ± (6.9) | 41.5 ± (6.9) | 41.1 ± (4.3) | 41.1 ± (6.0) | 39.0 ± (6.2) |
| Body fat (%) | 49.9 ± (5.7) | 49.3 ± (5.6) | NA | 48.5 ± (5.6) | 45.4 ± (8.6) |
| Kleiner fibrosis grade |  |  |  |  |  |
| F0 | 13 | 10 | 6 | 39 | 11 |
| F1 | 31 | 33 | 16 | 69 | 23 |
| F2 | 11 | 10 | 3 | 25 | 6 |
| F3 | 3 | 4 | 1 | 7 | 1 |
| F4 | 0 | 1 | 1 | 3 | 0 |
| SAF diagnosis |  |  |  |  |  |
| No MASLD | 12 | 16 | 13 | 39 | 22 |
| MASL | 29 | 30 | 11 | 74 | 14 |
| MASH | 17 | 12 | 3 | 30 | 5 |
| Steatosis |  |  |  |  |  |
| 0 | 12 | 16 | 13 | 39 | 22 |
| 1 | 22 | 27 | 12 | 60 | 11 |
| 2 | 17 | 9 | 1 | 28 | 7 |
| 3 | 7 | 6 | 1 | 16 | 1 |
| Hepatocellular ballooning |  |  |  |  |  |
| 0 | 41 | 46 | 24 | 112 | 36 |
| 1 | 16 | 12 | 3 | 29 | 4 |
| 2 | 1 | 0 | 0 | 2 | 1 |
| Lobular inflammation |  |  |  |  |  |
| 0 | 13 | 18 | 13 | 49 | 23 |
| 1 | 31 | 28 | 11 | 71 | 17 |
| 2 | 11 | 12 | 3 | 21 | 1 |
| 3 | 3 | 0 | 0 | 2 | 0 |

### Proteins involved in the innate immune system, extracellular matrix organization, and metabolism are differentially expressed in the MASLD liver

Robustly calculating differential expression between disease stages require complete data. However, MS-based proteomics is inherently prone to random missingness, and considering human heterogeneity we did not want to impute any missing values. Instead, we used ridge regression to prevent overfitting during statistical analysis of differential expression^15^. To improve model performance, we first performed an unsigned weighted gene co-expression network analysis (WGCNA)^16^ to limit feature size and increase feature similarity in the dataset. Analysis of differential for Kleiner fibrosis grade and SAF diagnosis resulted in 304 and 256 significantly differentially expressed proteins (DEPs) respectively. To improve statistical confidence and ensure biological relevance, we carried out any further analysis using only the 234 overlapping DEPs between Kleiner fibrosis grade and SAF diagnosis (Fig. 1b).

To verify the relationship between the DEPs and MASLD histopathology, we profiled the protein-wise relative expression after sorting patients based on increasing severity of MASLD histopathology. After hierarchical clustering of DEPs, we observed two clusters of proteins pertaining to being either up-or downregulated in expression (Fig. 1c). Using pathway analysis, we found the most significantly downregulated proteins to all originate from various pathways of metabolism, while the upregulated proteins belonged to pathways related to extracellular matrix (ECM) organization and immune response (Fig. 1d,e).

### Integration of public single-cell data links MASLD proteome changes to individual cell types

Although single-cell omics play a crucial role in deepening our understanding of disease mechanisms, single-cell proteomics has yet to become a routine methodology. To address this gap, we explored whether single-cell transcriptomic data from human liver could assist in deciphering our bulk proteomics data, aiming to provide insights into the potential cellular origins of the DEPs. To achieve this, we integrated three publicly available datasets^17–19^, which we annotated collectively based on cell type-specific gene expression markers and utilized the results to extrapolate a broad overview of the cell type-specific context of the DEPs.

Hierarchical clustering based on the predicted cell type-specific contribution towards the expression of the DEPs resulted in five clusters (Fig. 2a, Supplementary Fig. S3). As expected, the largest cluster, Cluster 1, consisted of DEPs predicted to be mainly expressed by the hepatocytes. We found the hepatocyte cluster to be the primary contributor of proteins associated with a decrease in metabolism, specifically in amino acid and fatty acid metabolism (Fig. 2b). Among the upregulated DEPs in hepatocytes, we found pathways of MET signaling. The MET receptor tyrosine kinase signals via classical mitogen-activated protein kinase (MAPK) signaling cascades but can, after binding to its ligand, hepatocyte growth factor (HGF), also transactivate other receptors, including the hyaluronan receptor CD44, ICAM1, and integrins^20^.

**Fig. 2:**
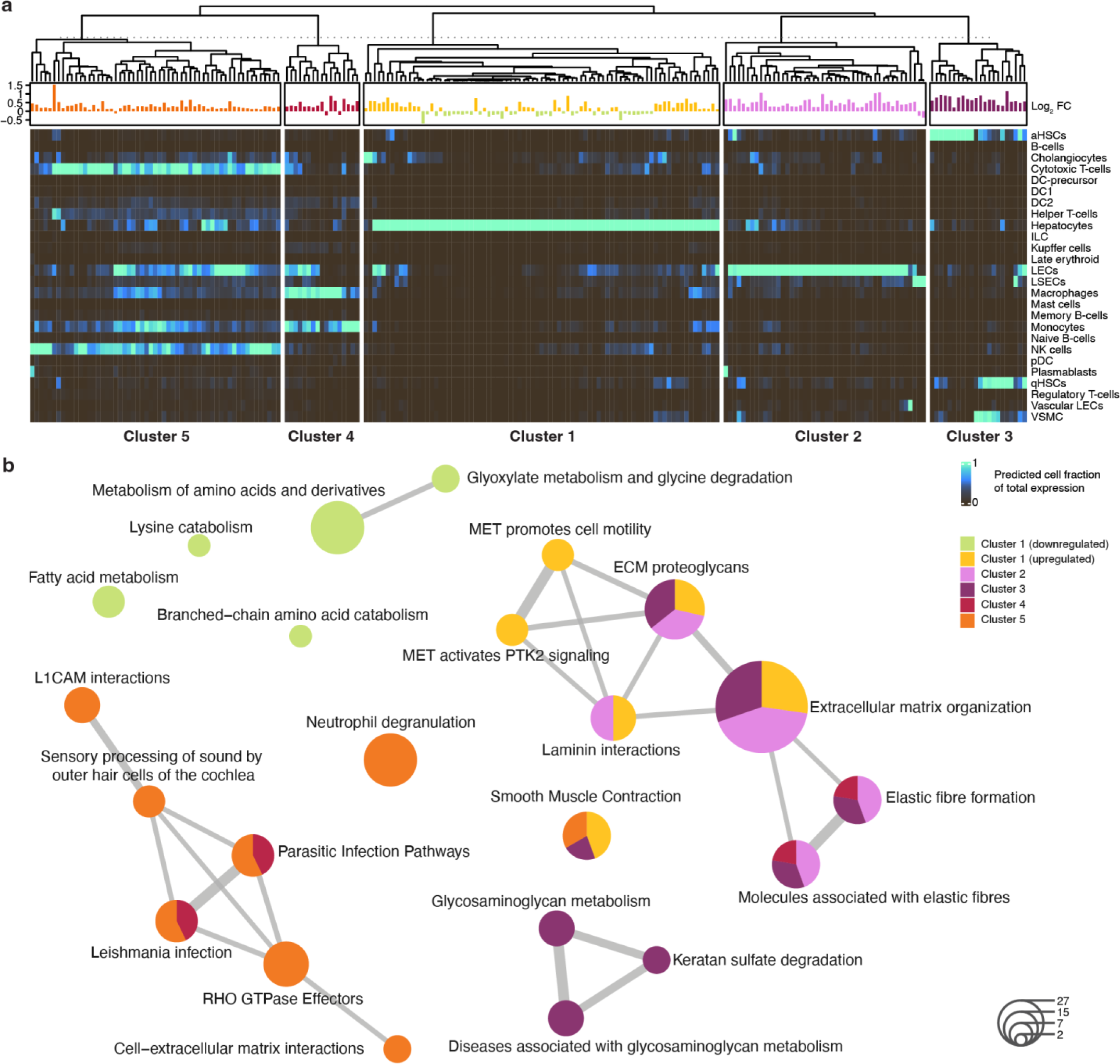
Public single cell data to unravel a cell type specific response in MASLD pathogenesis. **a,** Estimated cell type specific contribution for the total expression of our 234 significantly differentially expressed proteins in MASLD. Estimates are based on aggregated data from three single cell datasets, collectively annotated, count summarized by cell type, and then min-max normalized features. Each protein’s log_2_ fold change in expression shown on top (Supplementary Fig. S3). **b,** Comparison of enriched pathways for each cluster after calculating the pairwise term similarity matrix.

Cluster 2 consisted of DEPs expressed by liver endothelial cells (LECs), while Cluster 3 consisted of proteins mainly expressed in quiescent hepatic stellate cells (qHSCs), activated hepatic stellate cells (aHSCs), and vascular smooth muscle cells (VSMCs). High levels of VSMCs and aHSCs are major hallmarks of MASLD pathogenesis^21^. Pathway analysis showed that DEPs expressed by the main resident cell types – clusters 1, 2, and 3 – largely contribute to the organization of ECM. ECM organization is a convoluted process that involves many different regulatory networks. Proteoglycans are a major constituent of ECM and play an important role in cell signaling, e.g. by interacting with growth factors, cytokines, chemokines, and various pathogens. The architecture of the ECM provides many innate properties of tissues, and in MASLD pathogenesis, it is known to play a role in the advancement of fibrosis. In fact, liver stiffness measurements using elastiometry (i.e. Fibroscan) is today one of the best techniques in the non-invasive assessment of liver fibrosis^22^.

To further investigate some of the ECM-related sub-categorization, we explored the cell type-specific cellular processes (Fig. 2b). For example, we found laminin interactions primarily in hepatocytes and LECs. Laminins are known to commonly interact with integrins to control cell-to-ECM adhesion. In connection with this, in cluster 5 – mainly immune-related cytotoxic T-cells and natural killer (NK) cells – we found pathways of cell-ECM interactions as well as L1CAM interactions and RHO GTPase effectors. L1CAM interactions are known to be involved in cell-cell adhesion and RHO GTPases can regulate cell behavior, cell motility, and cytoskeleton organization^23^. This suggested a finding of proteins that were directly involved in the infiltration of immune cells into the tissue, and indeed, we found many of the proteins involved in the KEGG leukocyte transendothelial migration pathway to be upregulated during MASLD pathogenesis (Fig. 3a, Supplementary Fig. S4,5).

**Fig. 3:**
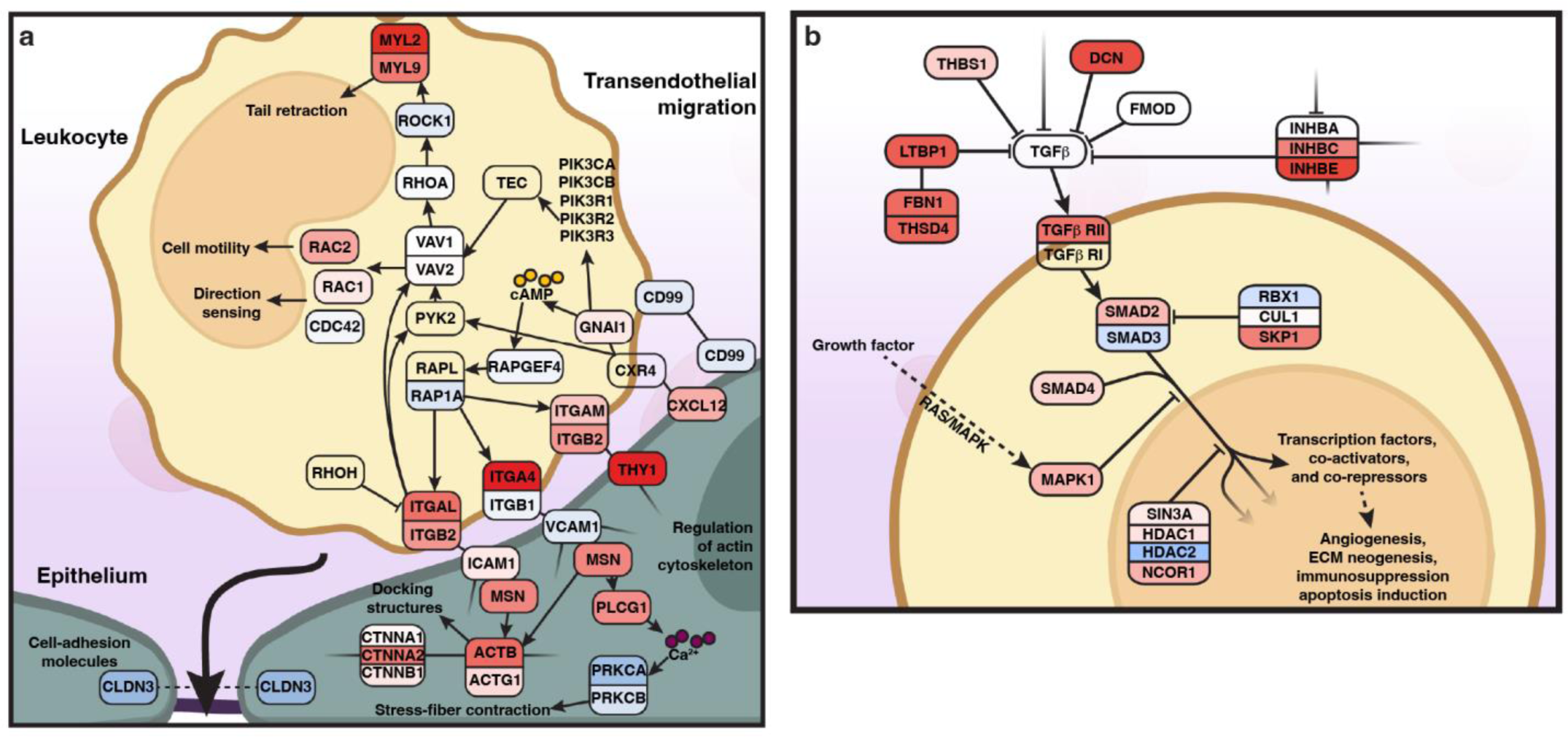
Depiction of signaling pathways related to MASLD pathogenesis. Proteins identified in **a,** transendothelial migration (hsa04670) and **b,** TGFβ signaling (hsa04350). Protein colors indicate change in expression where red is upregulated, blue is downregulated, white is no change, transparent is not identified. Details of full pathways and coloring schemes can be found in **a,** Supplementary Fig. S4,5 and **b,** Supplementary Fig. S6,7.

In cluster 5 – mainly immune cells – we found a large group of proteins involved in neutrophil degranulation. RHO GTPase-activated Rac2 has been shown to promote primary neutrophil degranulation of reactive oxygen species (ROS). Among the DEPs, we also found an upregulation of TP53I3 and GPX3 in the hepatocyte cluster, both of which are known to modulate ROS. Increased ROS is a known hallmark in MASLD pathogenesis, and both TP53I3^24^ and GPX3^25^ have already been suggested as a drug target and a potential biomarker respectively.

Another interesting observation related to glycosaminoglycan (GAG) metabolism, which is largely contributed by cluster 3 (HSCs). GAGs mainly consist of hyaluronan (HA; hyaluronic acid) and various sulfates (dermatan, keratan, heparan, and chondroitin) and act as a lubricating fluid in the ECM. HA is an anionic, nonsulfated glycosaminoglycan which has been suggested in multiple papers as a promising biomarker for MASLD^26^. Production of these GAGs can be activated by transforming growth factor-β (TGF-β) signaling. TGF-β signaling through SMAD2/SMAD3 in hepatocytes has been shown to accumulate free fatty acids, increase ROS production and mediate hepatocyte death^27^. Surprisingly, we also found an upregulation of elastic fibers in the LEC and HSC clusters, which, together with fibrillin microfibrils, can inhibit TGF-β signaling. When we checked for proteins related to TGF-β signaling, we found many of those to be differentially upregulated, but also many inhibitors of TGF-β (Fig. 3b, Supplementary Fig. S6,7). This could suggest a negative feedback mechanism or a mitigating response mechanism from nearby cells.

### Identification of transcriptional regulators that correlate with MASLD DEPs

To investigate the transcriptional regulators^28^ involved in MASLD pathogenesis, we performed a correlation analysis of all transcriptional regulators identified in our dataset to various patient characteristics (Fig. 4a, Supplementary Fig. S8). We then kept the transcriptional regulators with an accumulative correlation score above 0.45 across MASLD histopathology (SAF diagnose, Kleiner fibrosis grade, and NAFLD activity score) and performed hierarchical clustering of their correlation to the DEPs (Fig. 4b). Consolidating our earlier finding (Fig. 1 and 2), the expression patterns of the transcriptional regulators separated the DEPs into three major groups, group A consisting of pathways related to metabolism, group B relating to immune response, and group C relating to ECM organization (Fig. 4c-e). Further highlighting the robustness of the clustering results, we observed a positive correlation for SMAD2 with proteins involved in the immune response and a positive correlation between MYH11^29^ and ACTG2^30^, both being markers of VSMC. We also noted a positive correlation between the group of SMARC proteins with pathways of immune response, of which SMARCC1 has already been suggested as a putative prognostic marker for hepatocellular carcinoma (HCC) and targeted inhibition of SMARCC1 has been shown to reduce immune infiltration^31^.

**Fig. 4:**
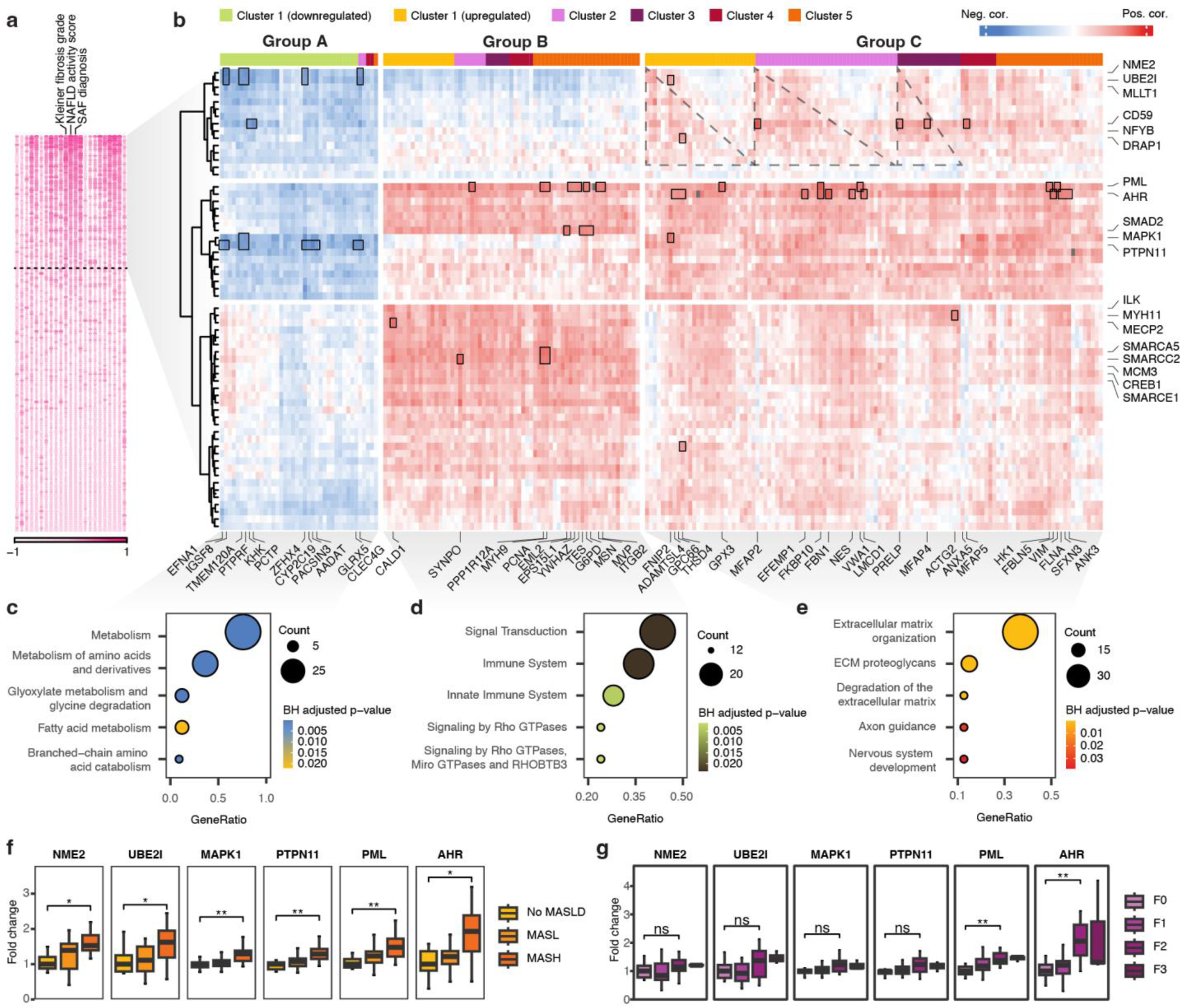
The role of transcriptional regulators in progression of MASLD. **a,** Correlation between all identified transcriptional regulators and patient characteristics, ordered by the sum of correlation to Kleiner fibrosis grade, NAFLD activity score, and SAF diagnose. Cutoff was arbitrarily selected. **b,** Hierarchical clustering of above cutoff transcriptional regulators with their correlation to the 234 significantly differentially expressed proteins of Kleiner fibrosis grade and MASLD. The columns (234 proteins) were first hierarchically clustered (Supplementary Fig. S9a), generating three clusters and then subsequently sorted within each of these three clusters by their cell type cluster (Fig. 2a). An asterisk indicates proteins with perfect correlation (found in both rows and columns). Squares indicate proteins with a higher correlation and were primarily chosen based on their correlation to the highest-ranking transcriptional regulators. The full figure containing all protein names can be found in Supplementary Fig. S9b. **c-e,** Pathway enrichment analysis based on the Reactome pathway database for proteins in each of the column-wise clusters. **f,g,** Fold change in protein expression for a selected subset of transcriptional regulators according to increasing **f,** SAF diagnosis and **g,** Kleiner fibrosis grade. Boxplot are shown in the style of Tukey (median, hinges: Q1 and Q3, and whiskers: 1.5x of IQRs from Q1 and Q3) and with symbols indicating statistical significance as calculated using Wilcoxon signed-rank test and BH correction method (ns p-value > 0.05, * p-value < 0.05, ** p-value < 0.01, *** p-value < 0.001, **** p-value < 0.0001).

Among the transcriptional regulators with overall highest positive correlation to the DEPs were aryl hydrocarbon receptor (AHR) and promyelocytic leukemia protein (PML) (Fig. 4f,g). AHR is a ligand-activated transcription factor that was recently suggested as a potential drug target in patients with MASLD^32^. However, AHR signaling pathways are not fully understood, and considering most AHR studies are in mouse models, we lack evidence for its potential involvement in human diseases. While PML followed a very similar profile to AHR, it has not yet been described in relation to MASLD and could potentially be a novel transcriptional regulator. PML acts as a tumor suppressor and is frequently found in nuclear bodies and nucleoplasm. However, emerging evidence suggests a multifaceted role that could make it important in the regulation of metabolism and immune response^33^. Interestingly, it has before been shown that PML can interact with UBE2I and SMAD2, both of which are among the top candidates of transcriptional regulators with the highest overall correlation to MASLD pathogenesis in our data (Fig. 4a).

As a major component of TGF-beta signaling, SMAD2 has already been linked to hepatic fibrosis, while UBE2I has mainly been associated with hepatocellular carcinoma (HCC)^34,35^. UBE2I is an E2 SUMO-conjugating enzyme and displayed a negative correlation with DEPs related to metabolism. Increased UBE2I expression has specifically been associated with autophagy processes and a poor prognosis for patients with HCC^34^. This negative correlation with pathways of metabolism could be a ripple effect from autophagy of dying hepatocytes, thus decreasing overall healthy hepatocyte function within the bulk of the analyzed tissue.

### Identification of MASLD-associated proteins in blood plasma

There is an urgent need to find robust, non-invasive biomarkers of MASLD. To test if any of the DEPs identified in the liver could also be found in the blood of the patients, as a result of cellular secretion or cell leakage, we analyzed 184 plasma samples from obese individuals by DIA MS-based proteomics (Table 1). First, we performed a correlation analysis of plasma proteins to MASLD histopathology and then ranked them based on their correlation to Kleiner fibrosis grade and SAF diagnosis (Fig. 5a,b). While there was an overlap between the top 10 most positively and negatively correlating proteins for Kleiner fibrosis grade and SAF diagnosis, they displayed unique signatures. For example, C7 and COLEC11 were specific in advancement of fibrosis but were not as important in determining MASLD. COLEC11 is known to be involved in activating the lectin complement pathway and generally recognizes pathogen cell surface sugars and apoptotic cells, guiding immune cell migration to clear microbes and heal injury^36^. C7 is also part of the complement system, being part of the membrane attack complex on target cells, and was differentially expressed in both significant and severe fibrosis (Fig. 5c). Another protein important for fibrosis was the negatively correlating IGF2 (Fig. 5a,d). A known transcriptional repressor of IGF2 expression is CTCF, which we also identified in our list of transcriptional regulators of MASLD pathogenesis (Supplementary Fig. S9). In relation to IGF2, we also found IGFALS, IGFBP3, IGFBP5, and IGF1 all being among the most downregulated in progression of liver fibrosis (Fig. 5a). IGFBP3 is known to interchangeably carry IGF1 and IGF2 in the circulation until they are released in the target tissue.

**Fig. 5:**
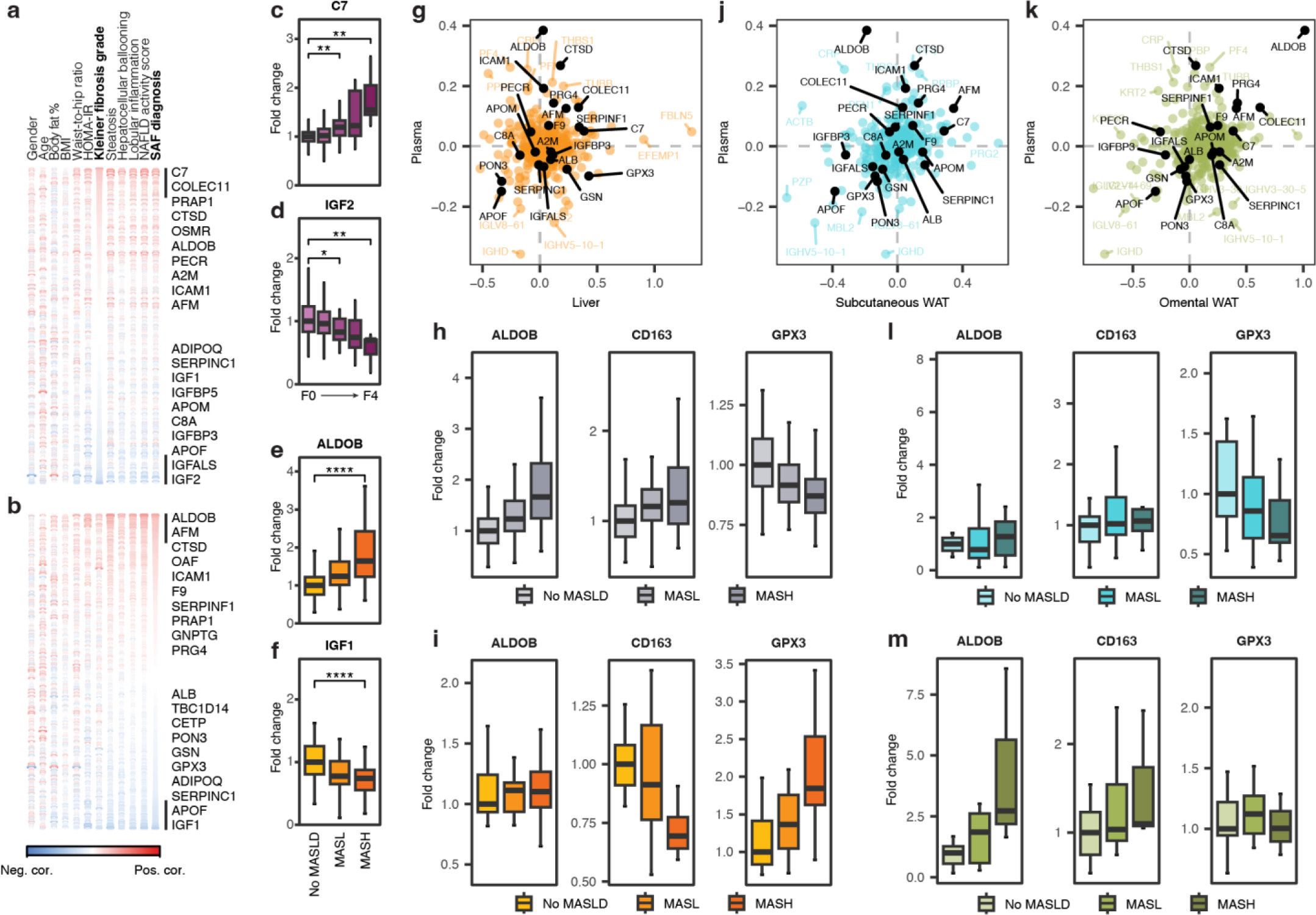
MASLD-related profiling of blood plasma. **a,b,** Correlation between all identified blood plasma proteins and patient characteristics, with top 10 most positively and negatively correlated proteins displayed according to increasing **a,** Kleiner fibrosis grade and **b,** SAF diagnose. **c-f,** Boxplots of proteins most positively and most negatively correlating with **c,d,** Kleiner fibrosis grade and **e,f,** SAF diagnose. **g,j,k,** Log2 of fold change protein expression during progression of MASLD in **g,** blood plasma vs liver, **j,** blood plasma vs scWAT, and **k,** blood plasma vs oWAT. **h,i,l,m,** Boxplots displaying relative fold change in expression for three proteins selected based on no correlation or a negative correlation between blood plasma and liver during progression of MASLD, for each of the four tissues; **h,** blood plasma; **i,** liver; **l,** scWAT; **m,** oWAT. Boxplot are shown in the style of Tukey (median, hinges: Q1 and Q3, and whiskers: 1.5x of IQRs from Q1 and Q3) and **c-f,** with symbols indicating statistical significance as calculated using Wilcoxon signed-rank test and BH correction method (ns p-value > 0.05, * p-value < 0.05, ** p-value < 0.01, *** p-value < 0.001, **** p-value < 0.0001).

When considering the top 10 most positively and negatively correlating proteins in relation to SAF diagnosis, we found ALDOB to be the most positively correlating candidate (Fig. 5b,e). ALDOB deficiency and ALDOB depletion, has been shown to result in increased intrahepatic accumulation of fatty acids and promoting hepatocellular carcinogenesis^37,38^. Interestingly, while IGF2 was most negatively correlating to Kleiner fibrosis grade specifically, for SAF diagnosis, it was IGF1 (Fig. 5b,f). In many cancers, including non-islet cell tumor hypoglycemia^39^, colorectal adenomas^40^ and cancers^41^, the molar ratios of IGF2/IGF1, IGF1/IGFBP3, and IGF2/IGFBP3 have already been used in diagnostics.

### WAT contributes to the plasma level differences of proteins associated with MASLD

Metabolic disorders are inherently complex, often emerging from interorgan crosstalk and multifaceted dysfunction across organs^42^. Among the blood plasma proteins correlating to MASLD histopathology (Fig. 5a,b), we find several that were absent in our liver-derived DEPs. This observation highlights the potential cross-organ dynamics related to MASLD. To explore this further, we first checked the expression differences for all blood plasma proteins against their expression profiles in the liver (Fig. 5g). Here, we found several examples of MASLD correlating proteins showing a discrepancy between blood plasma and liver. To highlight a few, GPX3 was downregulated in blood plasma while being upregulated in liver, and ALDOB was highly upregulated in blood plasma, while showing no changes in the liver (Fig. 5h,i). We also found an inverse relationship for CD163 where CD163 is downregulated in liver but upregulated in blood plasma, corroborating previous findings^43^.

Recognizing the pivotal role of adipose tissue in metabolic disorders^44^, we investigated if some of the changes in blood plasma correlating to MASLD pathogenesis that could not be explained by changes in the liver could instead be explained by adipose tissues. We performed DIA MS-based proteomic analysis of oWAT and scWAT from the same patient cohort. We then compared the change in expression for all blood plasma proteins against the changes observed in these adipose depots during MASLD pathogenesis (Fig. 5j,k). Indeed, for some of the MASLD-related proteins that could not be explained by changes in the liver proteome, we could correlate to adipose tissues. For example, ALDOB and CD163 were specifically upregulated in oWAT, matching the profiles observed in blood plasma for these proteins, while difference in expression for GPX3 was better matching the profiles observed in scWAT (Fig. 5l,m). These findings highlight the importance of considering multi-tissue interplay when studying complex diseases such as metabolic disorders.

### Predicting MASLD pathogenesis from blood plasma

Strategies relying on a single protein marker have not been very successful in diagnosis of MASLD-related pathologies. We therefore tested whether a proteomics-based model can be used as a “multi-protein panel marker” approach for predicting liver disease progression. For this purpose, we utilized the quantitation we had already attained using short-gradient DIA MS-proteomic analysis in plasma. We generated two 20-protein proteomics models, one designed to predict significant fibrosis (≥F2) and one for MASLD, based on their respective top 10 most positively and 10 most negatively correlating proteins for each histopathology (Fig. 5a,b). We also made a simplified version of the two 20-protein proteomics models consisting of only 5 proteins each – C7, ALDOB, ICAM1, CTSD, and IGF2 for significant fibrosis, and AFM, APOF, ALDOB, PRG4, and IGF1 for MASLD. We chose simple logistic regression for its ease of interpretability, and stress-tested the performance using 20x five-fold cross-validation. To evaluate model performance, we reported the area under the receiver-operating characteristic (AUROC), balanced accuracy, and F1 score (Fig. 6a-f). Our proteomics-based models, the 20-protein panel and the simplified 5-protein version, showed very similar results both performing better than existing frequently used models for predicting MASLD and fibrosis.

**Fig. 6:**
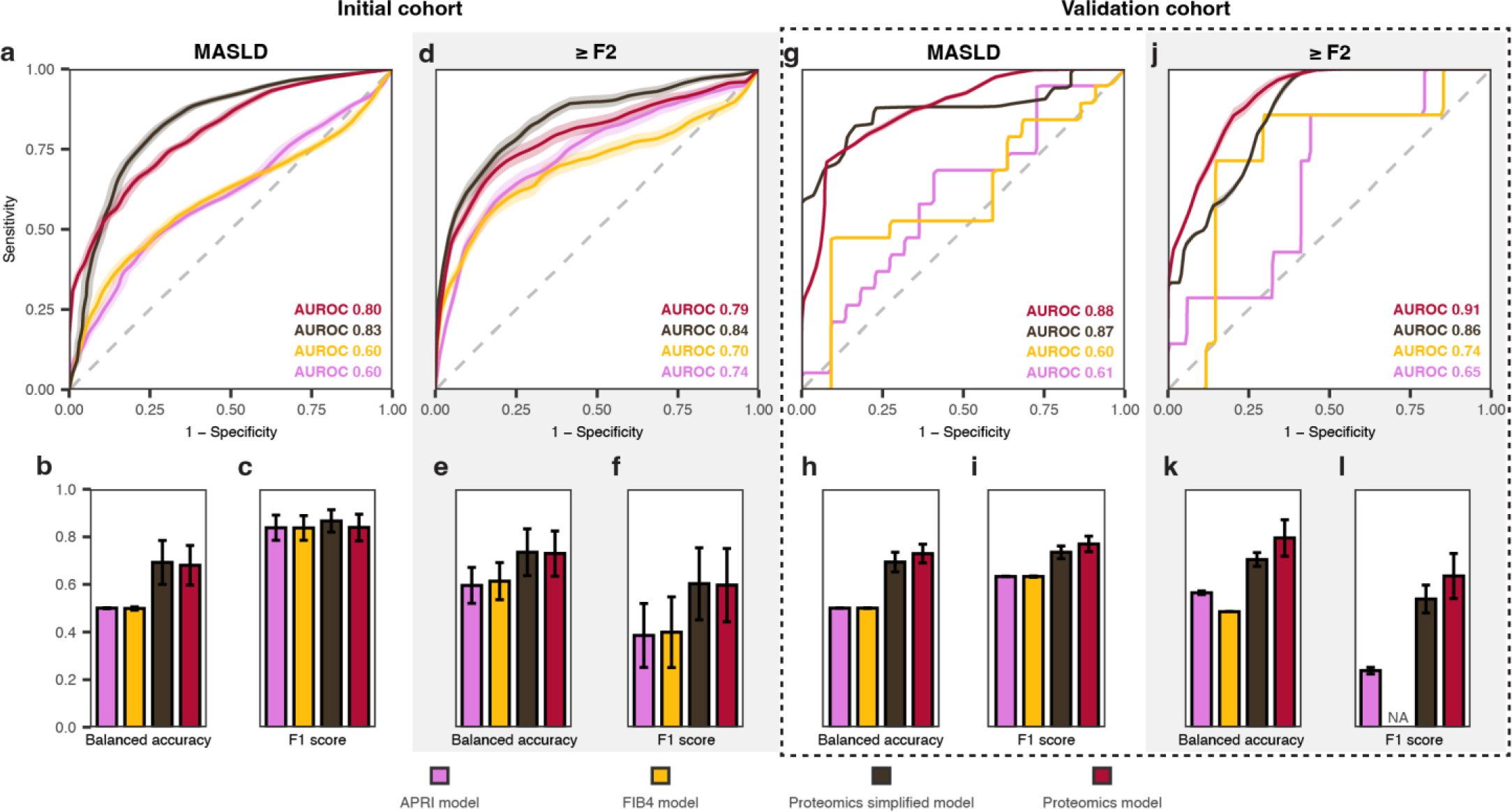
Performance evaluation of proteomics biomarkers for MASLD and ≥F2. Performance metrics for the logistic regression models after 20x five-fold cross validation (APRI, FIB4, 2x Proteomics models one for significant fibrosis (≥F2) and one for MASLD based on proteins from Fig. 6a and Fig. 6b respectively, and 2x Proteomics simplified models one for ≥F2 – C7, ALDOB, ICAM1, CTSD, and IGF2 – and one for MASLD – AFM, APOF, ALDOB, PRG4, and IGF1). **a-f,** Covers performance metrics of our initial cohort (n = 143), and **g-l,** covers our validation cohort (n = 41) consisting in follow-up blood samples. **a-c,** Model performance on predicting MASLD and **g-i,** again using the same model and model parameters on our validation cohort. **d-f,** Model performance on predicting ≥F2 and **j-l,** again using the same model and model parameters on our validation cohort. **a,d,g,j,** The receiver-operator characteristic (ROC) curve. **b,e,h,k,** The balanced accuracy. **c,f,i,l,** The F1 score. All plots show the mean across all one-hundred evaluations. Error bars in the bar plots show the standard deviation. Faded area in the ROC curves show the 95% confidence interval.

In addition to our initially screened plasma samples, we also received 2-year follow-up blood plasma from a subset of the cohort. From this subset, several patients had drastically reduced their body fat, with 13 out of 41 losing more than 10 BMI points due to bariatric surgery. Moreover, 31 out of the 41 patients had changes in their histopathology, of which 18 had improved their liver pathology scoring. We then used these 41 plasma samples as a validation cohort to evaluate the stability of our initial models’ performances in response to drastic phenotypic shifts within the initial cohort, We tested each individual model during the 20x five-fold cross validation on this cohort using the exact same parameters. Although the performance of the simplified 5-protein model was slightly reduced compared to the 20-protein panel, both proteomics-based models again overperformed the existing methods, which had a greater degree of instability (Fig. 6g-l). These results clearly substantiate the notion that such proteomics-based models relying on disease specific protein panels are indeed very robust and suitable strategy for characterization of complex diseases like MAFLD.

## Discussion

In this study, we took a proteomics-centric approach to explore MASLD in humans. We found an upregulation of immune response and extracellular matrix organization as key hallmarks of disease pathogenesis, accompanied by a downregulation of metabolism which could likely be a consequence of cellular reprogramming, loss of hepatocyte identity, and cell death^45^. We thoroughly explored the cell type-resolved signaling pathways and transcriptional regulators in MASLD pathogenesis, which we expect to be a valuable resource for future studies aiming to further investigate the cellular mechanisms in MASLD pathogenesis. Importantly, our findings reveal distinct expression patterns across liver, blood plasma, and WAT, highlighting the importance of considering interorgan crosstalk in understanding MASLD. Moreover, our disease-specific proteomics models proved to outperform existing diagnostic methods, offering a more accurate and reliable tool for predicting disease progression. Although rather speculative at this stage, we could envision similar strategies to replace the widely used single marker predictions in the clinics.

Previous studies have predominantly focused on a genomic and transcriptomic understanding of MASLD in humans. Here, we expanded the current understanding of MASLD from a proteomics perspective, while leveraging our findings for improving diagnostics. Interestingly, the difference in performance between our comprehensive 20-protein model and the streamlined 5-protein model was minimal. This suggests that good model performance hinges on its ability to capture essential disease aspects. With further optimization and the adoption of more sensitive quantification methods, we expect that the performance of our simplified models can be improved even further. By integrating comprehensive proteomic profiling across organs, we also uncovered potential crosstalk between liver and adipose tissues, specifically through proteins like ALDOB, CD163, and GPX3, whose differential expression suggests a nuanced mechanism of disease manifestation not previously studied.

Proteins comprise 75 % of all drug targets approved by the FDA and are the primary source of biomarkers^8^. By advancing our understanding of MASLD at a proteomic level, our work lays the foundation for improving diagnostic and therapeutic strategies. The correlation of adipose and plasma proteins with liver pathology not only aids in understanding MASLD mechanisms but also facilitates the development of targeted therapies. Our analysis highlighted proteins involved in the immune response and ECM organization, such as CD163 and GPX3. These proteins represent potential targets for new therapeutic interventions aimed at mitigating the inflammatory and fibrotic pathways in MASLD. Targeting GPX3, which is involved in the regulation of oxidative stress, could help in designing antioxidant therapies to reduce liver damage. Similarly, interventions that modulate CD163 activity could potentially limit the inflammatory responses, offering a therapeutic strategy to slow disease progression. Moreover, the identification of transcriptional regulators such as PML and AHR that correlate to GPX3 expression adds a layer of complexity and opportunity. Understanding these regulatory mechanisms further, can potentially lead to the precision therapies that target these pathways more effectively. Furthermore, our diagnostic proteomics-based biomarker models pave the way for non-invasive, cost-effective methods that could transform current practices by decreasing reliance on liver biopsies and enhancing early disease detection. They also provide a more nuanced insight for identifying MASLD. Leveraging differential expression patterns in key proteins, could potentially be utilized to predict disease trajectory in patients, which could enable more tailored interventions based on the projected course of the disease.

The strengths of this study include consistency in methodology across tissues and a well-described cohort with histological scoring for all samples. All samples across tissues were prepared using the same sample preparation procedure, processed in the same lab, same instrument setups, and using DIA-MS to vigorously fragment all eluting peptides. Using DIA-MS, we could use remains from the same biopsy that was used for histological scoring which would not have been feasible previously at such a depth. Moreover, conventional DDA would have suffered from high sample variance in human heterogeneity, introducing even more missingness and making the results more difficult to interpret. Being proteomics-based helps the translational potential of our findings in future diagnostic and therapeutic efforts.

Limitations to the study include potential artifacts created by human heterogeneity, cohort biases, and cohort size. To accommodate this, we took a conservative approach to the statistical analysis. While DIA-MS offers robust quantification and reproducibility, we did not consider post-translational modifications which play a huge role in cellular mechanisms and could be crucial in understanding MASLD pathogenesis. Given the discrepancy between RNA and protein expression levels, it should be expected that our cell type deconvolutions have a margin of error. Although our biomarker panel clearly correlates to disease progression, it does not necessarily reflect liver-specific mechanisms as already highlighted. MASLD is a multifaceted disease and investigating the direct mechanisms is outside the scope of this study. Analyzing multiple tissues adds complexity to the data interpretation, and in a human study like this we cannot derive any direct evidence for interorgan crosstalk.

Future research directions should employ longitudinal studies to monitor the dynamics throughout MASLD progression across relevant organs, and potentially explore even more organs than what we did here. Incorporating single-cell proteomics will also help delineate the cellular origins and functionalities of crucial proteins, providing mechanistic insights into therapeutic targets. Moreover, by integrating multiple omics we can develop a more comprehensive model that captures the intricate biological interplay underlying MASLD. Finally, by further optimizing our proteomics models and improving assay sensitivity, can advance the development of robust cost-effective non-invasive diagnostic tools, to reduce the need for liver biopsies.

In summary, we here provide exploratory proteomics-based study of MASLD, covering over 7,000 unique proteins in human liver biopsies from 58 obese patients with various degrees of liver disease progression. Using single-cell transcriptomic information to deconvolute our proteomics dataset, we linked many of the proteins associated with disease progression to individual cell types and the potential transcriptional regulators responsible for their expression changes in the disease states. The analyses uncovered more depth of the mechanisms related to the immune response and the extracellular matrix organization, including HSC differentiation, laminin and integrin interactions, RHO GTPase signaling, cell-ECM interactions, remodeling of LECs to accommodate transendothelial migration of immune cells, production of ROS, GAG metabolism, and regulation of TGF-β signaling. While being a liver-centric study, we nevertheless highlighted the multi-organ effects of MASLD in metabolic dysfunction trough quantitative proteomics analysis of scWAT and oWAT biopsies from the same 58 patient to a similar depth of ∼7000 unique proteins in each tissue. Last but not least, we utilized short-gradient DIA MS-proteomic approach for the analysis of 184 blood plasma samples and showed that our proteomics marker models outperform existing frequently used non-invasive methods in diagnosing MASLD. Overall, our findings provide rich and reliable resource, strengthening previous knowledge and bringing new insights that could help understanding normal liver function and MASLD pathology.

## Methods

### Patient cohort

**Study design**. Samples for proteomics were acquired from individuals participating in the PROMETHEUS study, a prospective interventional case-control study focusing on patients who undergo liver biopsy. PROMETHEUS is a single center study conducted in Denmark. **Inclusion and exclusion criteria.** Eligible participants must be in the age range 18 to 70 and have a BMI ≥ 35 kg m^-2^. The study excludes individuals with a limited life expectancy, excessive alcohol consumption (over 12 grams daily for women and 24 grams for men), any other chronic liver conditions, or usage of hepatotoxic drugs. **Ethics and registration**. The regional committee on health research ethics has granted approval for this study (S-20170210), which is registered at OPEN.rsyd.dk (OP-551, Odense Patient data Explorative Network) and at ClinicalTrial.gov (NCT03535142). Full ethical approval and informed written consent from participants were documented before initiation of the study. **Data collection**. Study data include anthropometrics and pharmacological treatment information. Data collection was carried out prospectively and managed using the REDCap (Research Electronic Data Capture) system hosted at OPEN.rsyd.dk, a secure, web-based platform designed to facilitate data capture for research purposes (https://www.sdu.dk/en/forskning/open).

### Histological staging

Liver biopsies were evaluated by a single trained radiologist (T.D.C), who was blinded to all other clinical data. The assessment adhered to the NASH Clinical Research Network (NAS-CRN) classification system^46^, with scoring for steatosis (0: <5%, 1: 5-33%, 2: 33-66%, 3: >66%), lobular inflammation (0: none, 1: <2 foci per 200x field, 2: 2-4 foci, 3: >4 foci), and hepatocellular ballooning (0: none, 1: moderate, 2: evident). These scores total to the NAS score (0-8). The FLIP (Fatty Liver Inhibition of Progression) algorithm and SAF (Steatosis, Activity, and Fibrosis) scoring system were used to further categorize patients based on the severity of their condition: no-MASLD (steatosis <1), MASL (steatosis ≥1), and MASH (steatosis ≥1, lobular inflammation ≥1, and hepatocellular ballooning ≥1)^14^. Fibrosis was staged using the Kleiner classification system, indicating the extent of tissue scarring (F0: none, F1: either perisinusoidal or portal/periportal fibrosis only, F2: both perisinusoidal and portal/periportal fibrosis, F3: bridging fibrosis, F4: cirrhosis)^46^.

### Sample acquisition

Liver biopsies, WAT biopsies, and blood samples were collected at the Department of Gastroenterology and Hepatology, Southwest Jutland Hospital in Esbjerg, Denmark. Patients were overnight fasting prior to sampling. Blood samples were collected in the morning prior to liver and WAT biopsies by specially trained hospital lab staff. Two designated lab techs handled blood sampling. Standard biochemical testing was done on the Cobas6000 (Roche). Blood components for the ATLAS biobank were frozen in a - 80 °C freezer immediately after aliquotation to secondary tubes. WAT was obtained immediately before the liver biopsy and was, as such, collected during the liver biopsy procedure through a 2 cm incision in the liver biopsy needle insertion area. Approximately 2×3 cm WAT was removed with a pair of Metzenbaum dissection scissors, cleaned in a sterile saline solution, frozen in liquid nitrogen, and then kept at −80 °C. Percutaneous liver biopsies from right liver lobe were acquired under sterile conditions using a 16-18G Menghini suction needle (Hepafix, Braun, Germany). The liver biopsy was cleaned in a sterile saline solution. At least 1.5 cm was sent for histologic evaluation, and the rest was used for research analysis. Liver biopsies for histologic staging (minimum 1.5 cm) were immediately stored in 4% formalin and embedded in paraffin. All biopsies for proteomics were immediately snap frozen in liquid nitrogen and then stored at −80 °C. Additional data, such as anthropometrics, were collected on the same day as biopsies and blood sampling.

### Protein purification and digest of biopsies

For quantitative proteomics, biopsies were sliced in two and homogenized in denaturing buffers (either 8 M guanidine hydrochloride in 25 mM ammonium bicarbonate or 5 % SDS) by 3-10 seconds of sonication (20 % intensity) and heated for 5 mins at 95 °C. WAT biopsies had an additional centrifugation step at 13,000 rpm for 10 mins to separate and avoid carry-on of superfluous lipids into the following steps. Pierce^TM^ BCA Protein Assay Kit (Thermo Fisher Scientific) was used to measure protein concentrations, and 10 μg of protein (5 μg from each lysate) was combined in a 1:1 volume before diluting 4x with water for a final concentration of 1 M guanidine hydrochloride (and 0.625 % SDS). For protein aggregation capture on microparticles ^47^ we used 70 % (v/v) acetonitrile (ACN), 40 μg of magnetic HILIC beads (ReSyn Biosciences (Pty) Ltd.) and 30 min incubation at room temperature with agitation on. Bead-bound protein aggregates were retained using a magnet while washing with pure ACN, followed by washing with 70 % (v/v) ethanol. Bead-bound protein aggregates were then resuspended in 100 μL of 50 mM ammonium bicarbonate, reduced with 2 mM DTT for 30 min, and alkylated with 11 mM chloracetamide for 30 min in the dark. Proteins were proteolytically digested using 1:200 of LysC (Wako) for 1 hour at 37 °C, followed by 1:100 of trypsin (Promega) overnight at 37 °C. Samples were acidified (pH 2.0-2.5) using trifluoroacetic acid, loaded onto in-house prepared C18 StageTips, desalted, and eluted again using 45 μL of 60 % ACN in 0.5 % (v/v) acetic acid. Finally, eluates were vacuum-dried to 2-3 μL in a speed-vac and re-diluted to ∼10 μL using 0.5 % (v/v) acetic acid.

### Preparation of blood plasma

Blood plasma was first diluted in 50 mM ammonium bicarbonate to a final concentration of 1 μg/μL. Next, we prepared 10 μg of protein in a 1:10 dilution of 50 mM ammonium bicarbonate, by reduction, alkylation, protein digest, and desalting on StageTips, as described above.

### Data-independent acquisition mass spectrometry

∼800 ng of digested and purified peptide mixture was loaded onto an LC-MS/MS system consisting of an EASY-nLC 1000 nanoflow liquid chromatograph (Thermo Fisher Scientific) coupled with either a Q Exactive HF-X mass spectrometer (Thermo Fisher Scientific) or an Orbitrap Exploris 480 mass spectrometer (Thermo Fisher Scientific)^48^. For reverse-phase chromatography we used a 20-cm analytical column with an inner diameter of 75 μm packed in-house with ReproSil Pur C18-AQ 1.9 μm resin (Dr Maisch GmbH), paired with an ACN/water (0.5 % acetic acid) solvent system at a flow rate of 0.25 μL min^-1^. For peptide separation of biopsy proteomes, we used a step-gradient from 3 to 8 % ACN for 7 min, followed by a slow gradient to 35 % ACN for 86 min, a ramp to 45 % ACN for 15 min and a plateau of 95 % ACN for 5 min. For peptide separation of plasma proteomes, we used a step-gradient from 3 to 8 % ACN for 1:22 min:sec, followed by a slow gradient to 35 % ACN for 16:43 min:sec, a ramp to 45 % ACN for 2:55 min:sec and a plateau of 95 % ACN for 5 min. The mass spectrometers were operated by sequential window acquisition of all theoretical fragment ion spectra^48^. For MS1, we used a 350 to 1400 m/z survey scan with a resolution of 120,000, a maximum ion injection time of 45 ms, and a normalized automatic gain control target of 300 %. This was followed MS2, fragmentation of precursor ions by higher-collisional dissociation at a collision energy of 28 % for each 13 m/z isolation window, and a 1 m/z overlap between sliding windows to sequentially cover the 360.5-1000.5 m/z range by 50 scans. For MS2, resolution was set to 30,000 and the maximum ion injection time to 54 ms. DIA raw files were analyzed in Spectronaut^TM^ version 18.1 (Biognosys) using default settings, and our in-house generated spectral library for WAT and liver, and DirectDIA (library free) for plasma. The report table containing protein identifications and quantitative values were kept for subsequent data analysis in R.

### In-house generated spectral library

Briefly, our in-house spectral library was generated using Pulsar in Spectronaut^TM^ (Biognosys) from data files of previous analyses of human liver using high pH fractionation and DDA. In total, our spectral library for liver contains 8678 unique proteins from 139,207 stripped peptide sequences (199,022 modified peptides sequences). While our spectral library for WAT contains 7672 unique proteins from 113,270 stripped peptide sequences (127,551 modified peptides sequences).

### Public single cell RNA-seq data integration and annotation

For deconvolution, three human single-cell RNA-sequencing (scRNA-seq) datasets were retrieved from public repositories GSE115469^17^, GSE136103^18^, and GSE158723^19^. Seurat (v.4.0.3)^49^ was used to process each dataset. Low quality cells were excluded (200< n <3,000 genes, mitochondrial gene contributions <20%). Moreover, genes expressed in fewer than 50 cells were excluded to remove zero count genes. Following cell exclusion, normalization, scaling, and dimensional reduction were performed. DoubletFinder (v.2.0.3)^50^ was employed to predict and remove doublets. For dataset integration, the processed datasets were merged, and the principal component embeddings were corrected using Harmony (v.0.1.0)^51^. CellTypist (v.0.1.4)^52^ was employed for automated cell type annotation of the complete dataset (trained model reference = Immune All Low). Manual correction of cell type annotations was performed to increase annotation resolution for hepatocytes, cholangiocytes, liver sinusoidal endothelial cells, liver endothelial cells, aHSCs, qHSCs, and VSMCs.

### Data analysis

**Software.** Analysis was performed using the open-source statistics software R, version 4.3.1 (2023-06-16) -- “Beagle Scouts”. **Analysis of differential expression.** We first performed quantile normalization of the summarized protein quantifications after log2 transformation to achieve normal distribution. We filtered proteins missing in more than half of all the samples and then performed unsigned network topology analysis, and determined the optimal power, based on the scale free topology model fit and the mean connectivity (Supplementary Fig. S1a). We used a power of 11 to calculate unsigned adjacency, which was then used to calculate the topological similarity and corresponding dissimilarity matrices, for hclust-based dendrogram and gene module calculation, using the flashClust, cutreeDynamic, and labels2colors functions (Supplementary Fig. S1b). We used the WGCNA R package^16^ to calculate module eigengenes and correlate eigengenes to patient characteristics by Pearson correlation (Supplementary Fig. S1c). Proteins from Black and Greenyellow gene modules, 348 proteins in total, was kept for further data analysis. Proteins were fitted using robust regression (MSqRob2 integration of the MASS rlm-function). Test statistics and Benjamini-Hochberg (BH) correction of p-values for all contrasts of Kleiner fibrosis grade and SAF diagnosis was calculated using the topFeatures-function in MSqRob2^15^. The overlap between statistically significant proteins (BH-adjusted p-value < 0.05) for Kleiner fibrosis grade and SAF diagnosis resulted in a total of 234 proteins. **Pathway enrichment analyses**. We used the ReactomePA R package^53^. Analysis was performed using the enrichPathway function and BH correction. **Estimation of cell type contribution**. For our deconvolutions, we did a count summarization for each cell type, followed by feature-wise min-max normalization. **Heatmap sorting**. Specifically for Fig 4b, we first performed hierarchical clustering, and then subsequently sorted each of the three major protein groups according to their cell type clusters for easier interpretation. The original heatmap can be found in Supplementary Fig. S9. **Statistical significance**. For analysis of significance for individual proteins shown in the barplots we performed a Wilcoxon signed-rank test and BH correction method. **Logistic regression models**. We performed 20x five-fold cross-validation logistic regression using random seeds. For each experiment we calculated the AUROC, balanced accuracy, and F1 score and then reported the mean and standard deviation. **Data visualization**. All data visualization were made using ggplot2^54^, enrichplot^55^, corrplot^56^, VennDiagram^57^, ComplexHeatmap^58^, Pathview^59^, and Adobe Illustrator.

## Data availability

The mass spectrometry proteomics data have been deposited to the ProteomeXchange Consortium via the PRIDE^60^ partner repository with the dataset identifier PXD051911.

## Code availability

All bioinformatic analyses were performed in the open-source statistics software R, version 4.3.1 (2023-06-16) -- “Beagle Scouts”. The packages and general details of the analysis is discussed briefly in the Methods. R scripts are available from the corresponding author upon reasonable request.

## Acknowledgements

This research received funding from the Danish National Research Foundation (grant number DNRF141) awarded to the Center for Functional Genomics and Tissue Plasticity (ATLAS) The proteomics work was supported by the INTEGRA mass spectrometry research infrastructure established at SDU by a generous grant from the Novo Nordisk Foundation (#NNF20OC0061575).

## Author contributions

Conceptualization: F.B. and B.B.; Proteomics laboratory work: F.B., V.A., M.T., and P.H.; Clinical work: C.W.W. and M.M.L.; Cell type deconvolutions: M.K.T and F.T.L.; Data nalysis and figures: F.B.; Supervised the study: B.B., M.M.L., A.K., K.R, and S.M.; Manuscript preparation: F.B.; All authors provided critical feedback to the manuscript and approved its final version.

