## Supplementary Figures S1-S9 for "Deep Proteome Profiling of Metabolic Dysfunction-Associated Steatotic Liver Disease"

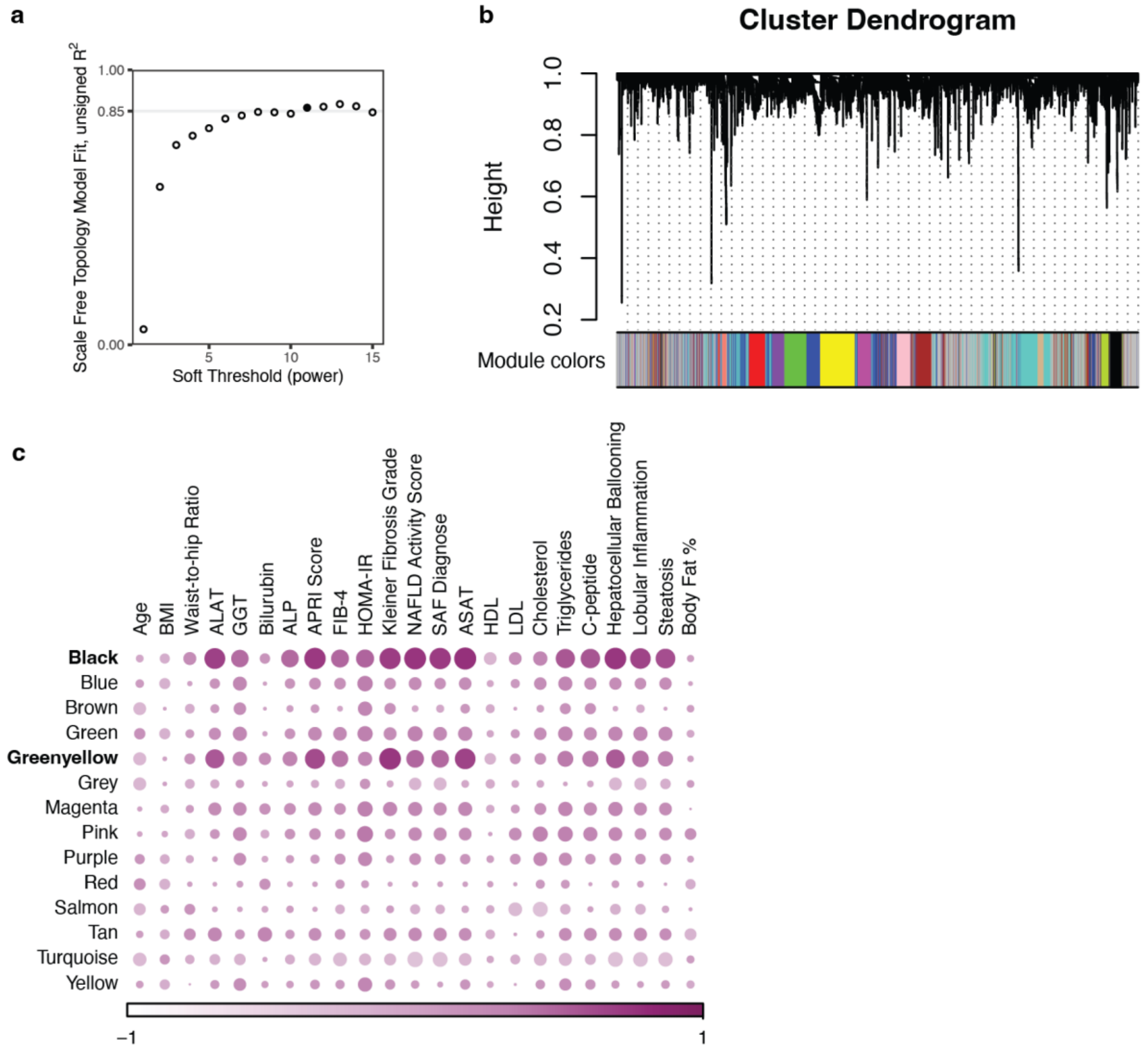


**Fig. S1 | Supporting figure for 'Methods' on 'Analysis of differential expression' regarding WGCNA. a,** Scale free topology model fit, a soft threshold power of 11 reaches >0.85 cutoff (black fill). **b,** Cluster dendrogram of hierarchical clustering. **c,** Correlation between eigenvectors of the gene modules and various patient characteristics.

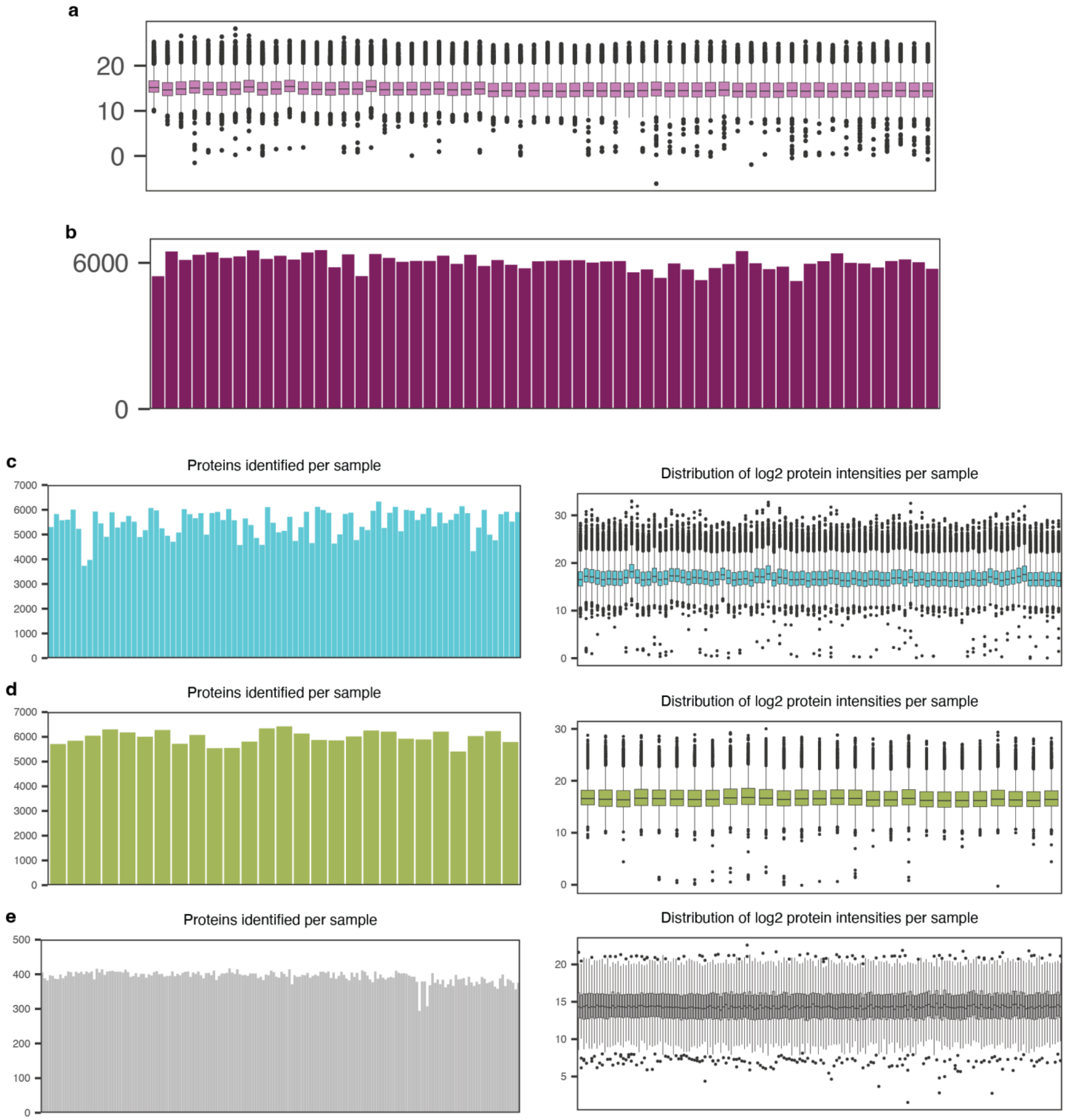


**Fig. S2 | Supporting figure to 'Study design and data collection' regarding proteomics sample variability.** Overview for liver samples, **a,** distribution of quantitation after log2 and **b,** number of unique proteins. Left shows the number of unique proteins identified and right shows the distribution of quantitation after log2 in **c,** scWAT, **d,** oWAT, and **e,** blood plasma.

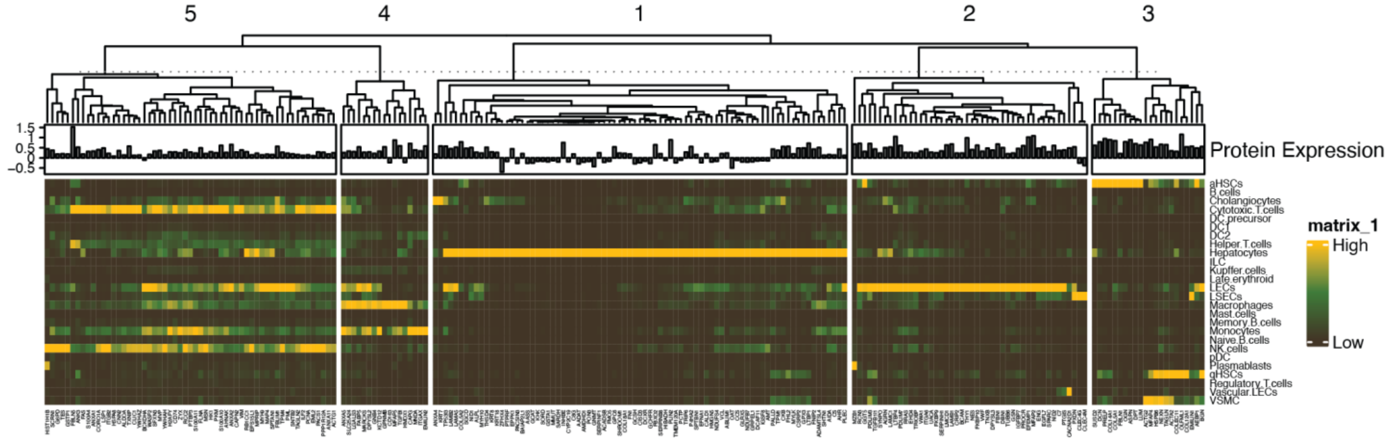


**Fig. S3 | Supporting figure for hierarchical clustering of cell type-specific contribution of the DEPs.** Same as Fig. 2a, but also showing protein names, available for transparency of clustering and to serve as a resource for future studies.

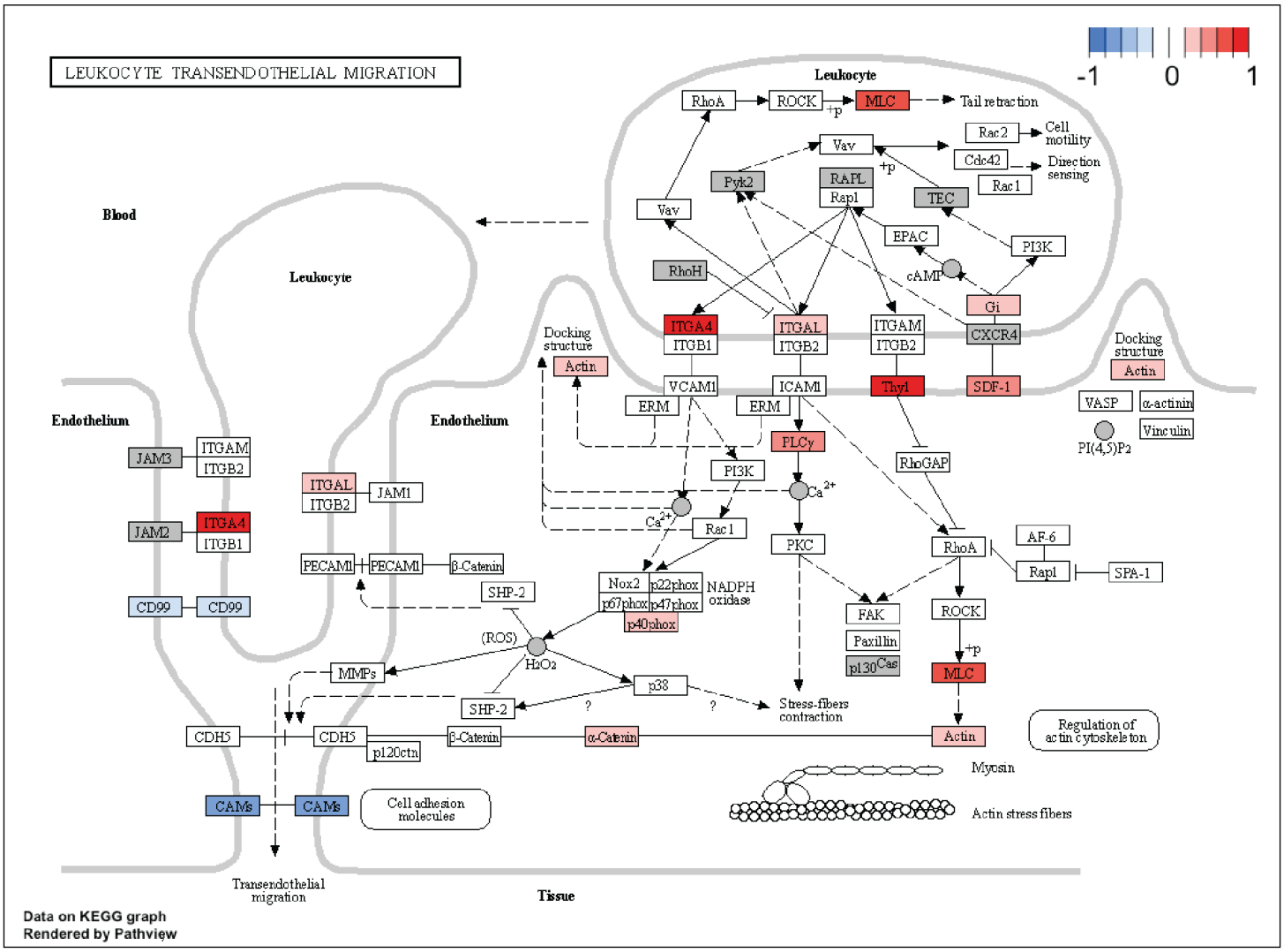


**Fig. S4 | Supporting figure for the depiction of transendothelial migration.** Served as the foundation for the illustration (Fig. 3).

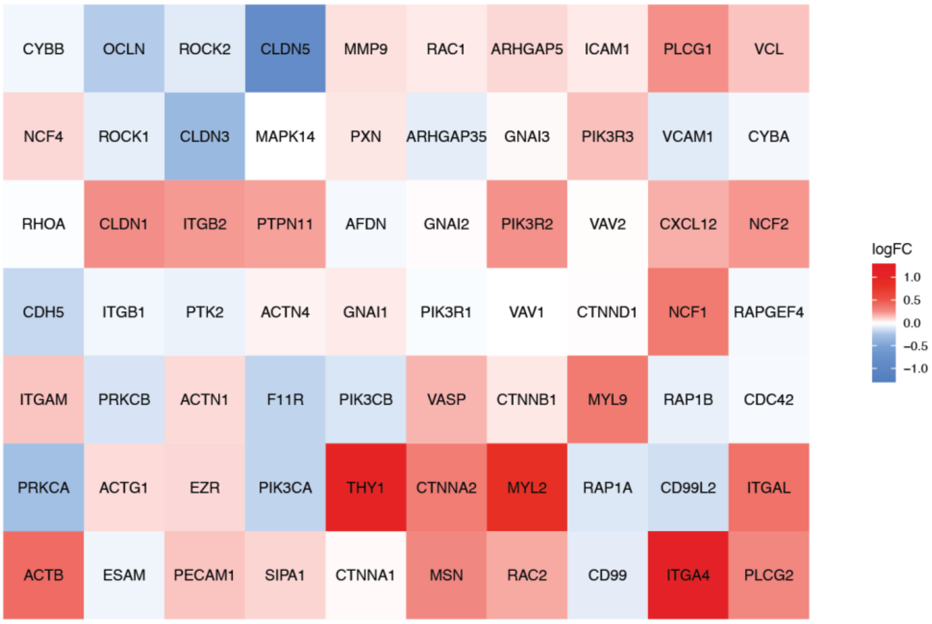


**Fig. S5 | Supporting figure for the depiction of transendothelial migration.** Used for the coloring in Fig.3.

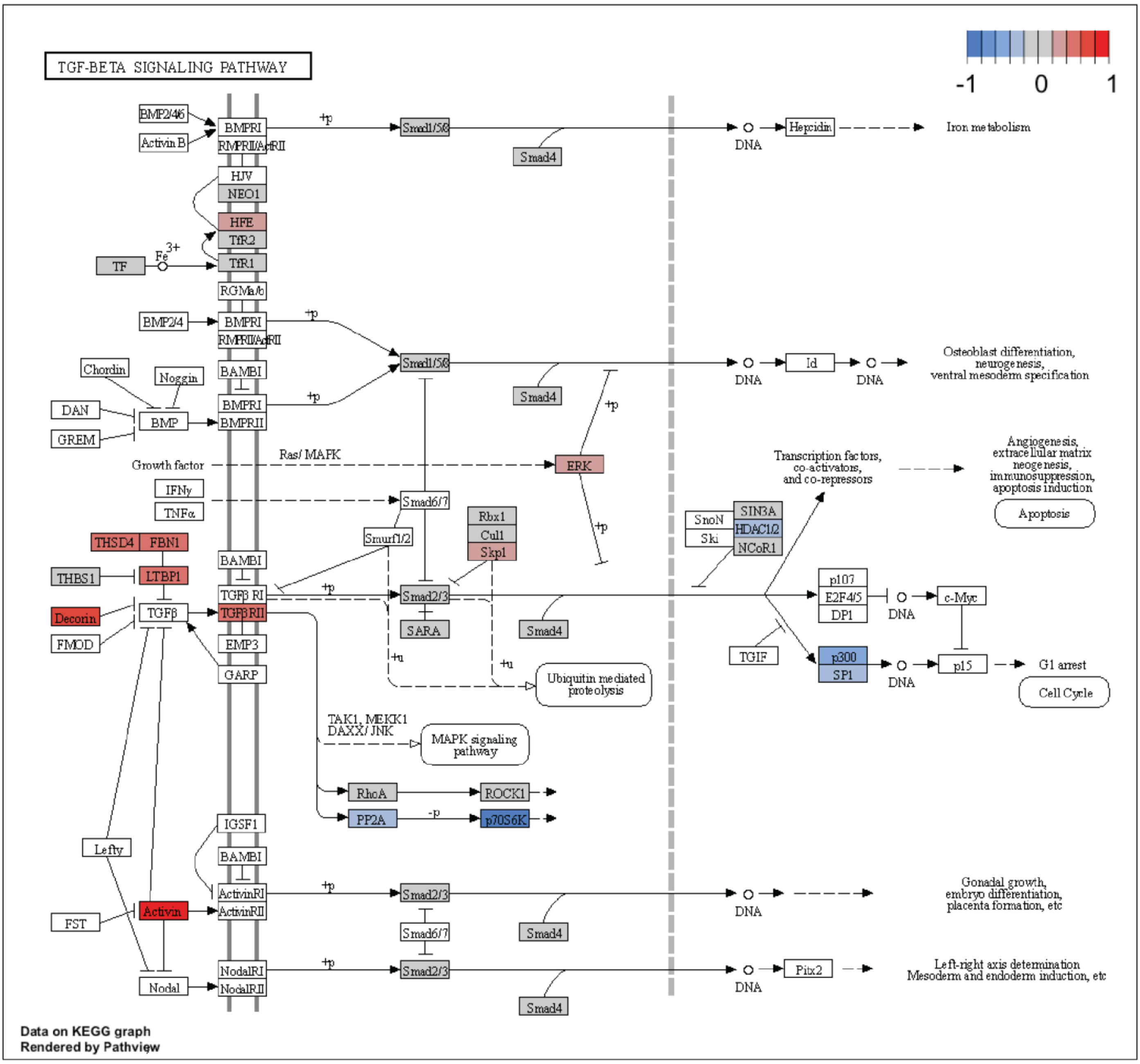


**Fig. S6 | Supporting figure for the depiction of TGFβ signaling.** Served as the foundation for the illustration (Fig. 3).

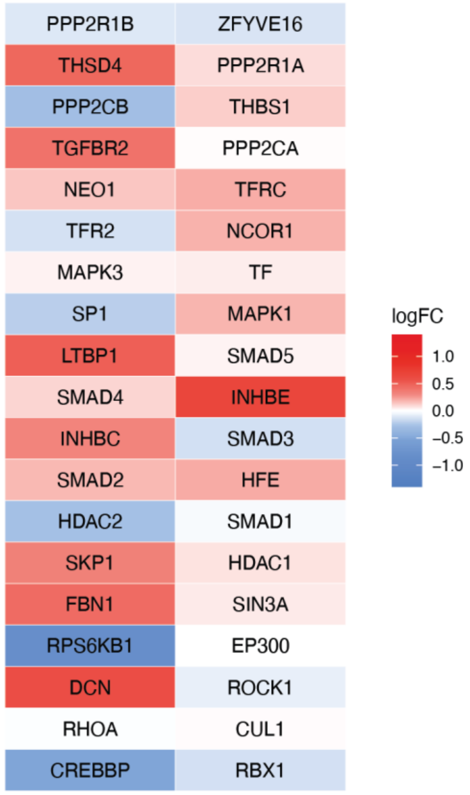


**Fig. S7 | Supporting figure for the depiction of TGFβ signaling.** Used for the coloring in Fig.3.

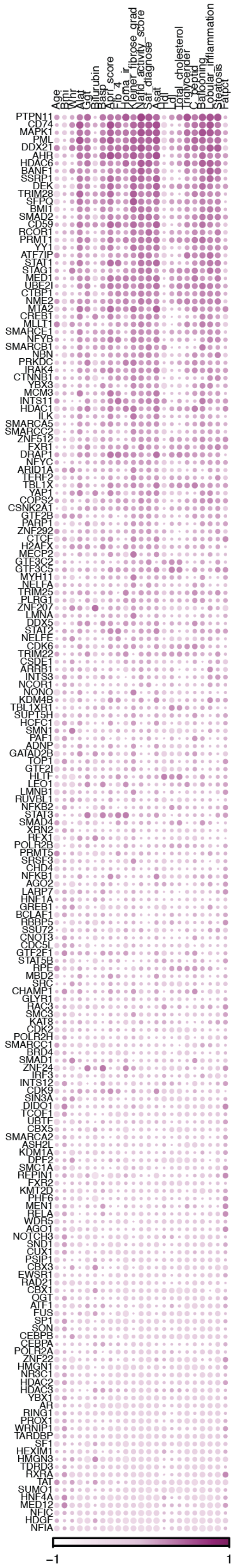


**Fig. S8 | Supporting figure for the correlation analysis between all transcriptional regulators and various patient characteristics.**

Same as Fig. 5a, but also showing names of the transcriptional regulators, available for transparency and as a resource for future studies.

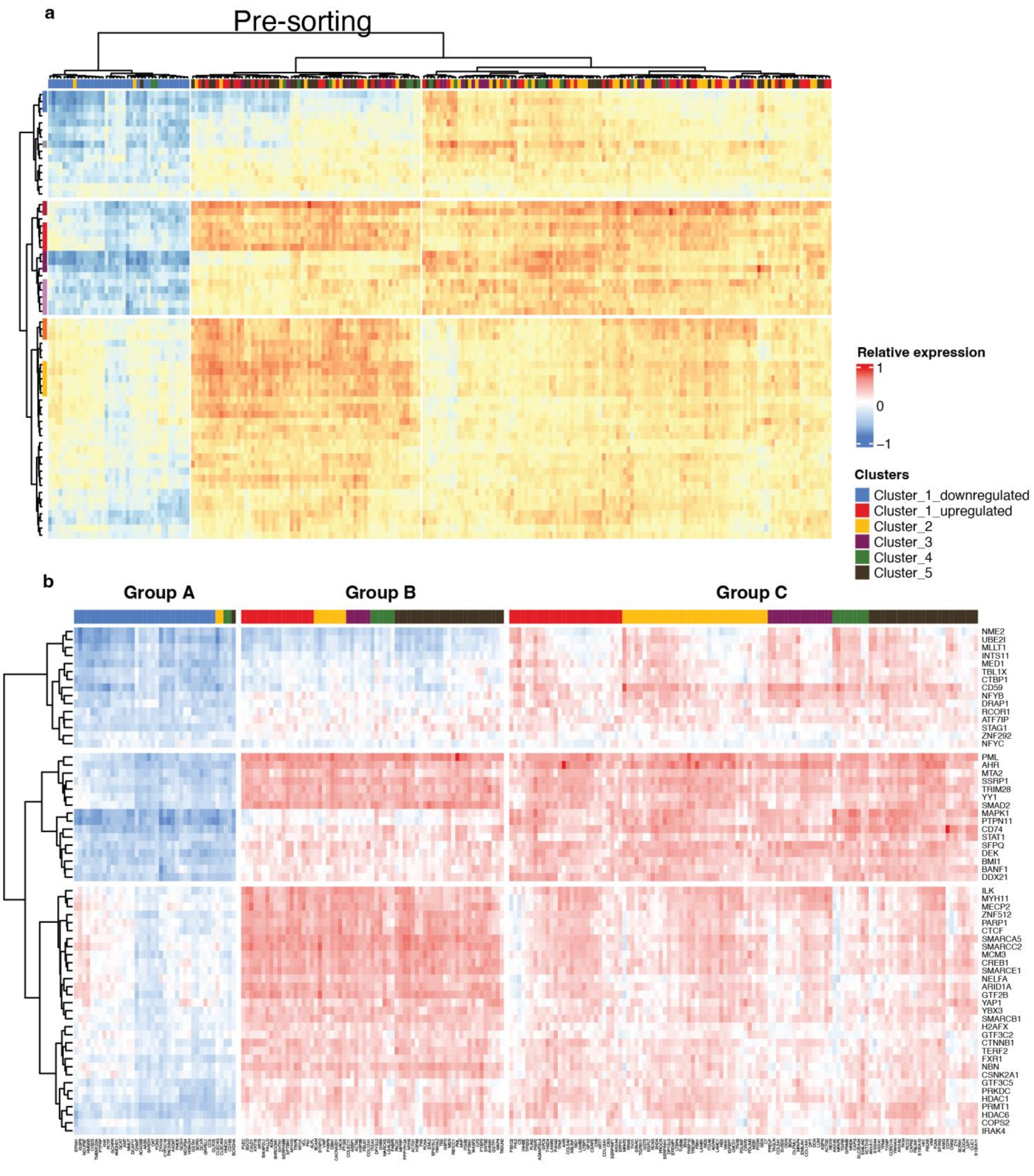


**Fig. S9 | Supporting figure to the correlation between MASLD DEPs and transcriptional regulators.** The hierarchical clustering **a,** before sorting and **b,** after sorting DEPs according to their cell-type clusters within each of the three group, displaying all names of DEPs and transcriptional regulators.
